# Fine-tuning Large Language Models for Rare Disease Concept Normalization

**DOI:** 10.1101/2023.12.28.573586

**Authors:** Andy Wang, Cong Liu, Jingye Yang, Chunhua Weng

## Abstract

**Objective:** We aim to develop a novel method for rare disease concept normalization by fine-tuning Llama 2, an open-source large language model (LLM), using a domain-specific corpus sourced from the Human Phenotype Ontology (HPO).

**Methods:** We developed an in-house template-based script to generate two corpora for fine-tuning. The first (NAME) contains standardized HPO names, sourced from the HPO vocabularies, along with their corresponding identifiers. The second (NAME+SYN) includes HPO names and half of the concept’s synonyms as well as identifiers. Subsequently, we fine-tuned Llama2 (Llama2-7B) for each sentence set and conducted an evaluation using a range of sentence prompts and various phenotype terms.

**Results:** When the phenotype terms for normalization were included in the fine-tuning corpora, both models demonstrated nearly perfect performance, averaging over 99% accuracy. In comparison, ChatGPT-3.5 has only ∼20% accuracy in identifying HPO IDs for phenotype terms. When single-character typos were introduced in the phenotype terms, the accuracy of NAME and NAME+SYN is 10.2% and 36.1%, respectively, but increases to 61.8% (NAME+SYN) with additional typo-specific fine-tuning. For terms sourced from HPO vocabularies as unseen synonyms, the NAME model achieved 11.2% accuracy, while the NAME+SYN model achieved 92.7% accuracy.

**Conclusion:** Our fine-tuned models demonstrate ability to normalize phenotype terms unseen in the fine-tuning corpus, including misspellings, synonyms, terms from other ontologies, and laymen’s terms. Our approach provides a solution for the use of LLM to identify named medical entities from the clinical narratives, while successfully normalizing them to standard concepts in a controlled vocabulary.

## BACKGROUND

Often individually rare but collectively common, rare diseases exhibit complex genetic heterogeneity and phenotypic manifestations. Both their phenotypes and disease concepts have heterogeneous references or documentation. For instance, one doctor might note “hearing loss” for a patient while another doctor might characterize the same patient as “having difficulty hearing.” A lack of standards-based concept normalization can lead to underdiagnosis, misdiagnosis or mistreatment [1]. Adopting a standard clinical vocabulary would make patient data more accessible to interpret and share, and further improve patient outcomes [2]. More importantly, standardized clinical concepts are crucial for efficiently and accurately studying trends and outcomes based on data aggregated from a large number of patients. The heterogeneous clinical data combined with insufficient comprehensive rare disease knowledge only hampers and reduces the efficiency of rare disease research, which further may compromise the quality and reliability of clinical findings [3-6]. The adoption of a standardized vocabulary holds the promise of simplifying clinical data significantly, allowing researchers to easily compare and analyze data across multiple medical settings and databases, accelerating medical research [7-9].

While standardized vocabularies for rare diseases, such as the Human Phenotype Ontology (HPO) have been established, their integration into clinical settings remains infrequent [10,11] and rare disease documentation in clinical settings remains unstandardized. Consequently, researchers often find themselves manually phenotyping patients using standardized vocabularies or, on a larger scale, employing Natural Language Processing (NLP) to recognize these standardized concepts from clinical narratives. In the latter scenario, the traditional approach typically involves a two-step process: concept recognition and concept normalization (such as translating “Gastroparesis” to “HP:0002578”). For instance, Doc2Hpo has utilized traditional NLP parsers, such as MetaMap [12], to identify terms within clinical text and then employ an indexing-based methodology to normalize these terms to standardized HPO concepts [13,14].

While NLP offers a solution, the effectiveness of traditional two-step processes is hindered when faced with slight modifications in clinical data and an inability to adapt to varying textual contexts. For instance, terms like “hearing loss” and “difficulty hearing” are not explicitly indexed as HPO names or synonyms. Therefore, traditional indexing-based normalization approaches may struggle to correlate them with the standardized HPO concept of “Hearing Impairment” (HP:0000365). Consequently, there is a pressing need for the development of more adaptable NLP tools to address the challenges associated with clinical concept normalization. Recent advancements in Large Language Models (LLMs), such as ChatGPT come with incredible contextual interpretability abilities backed by a myriad of knowledge. These models are characterized by their deep neural network architecture, typically consisting of Transformer-based models with billions of parameters [15]. LLMs are trained on vast amounts of text data, such as books, articles, and websites, using unsupervised learning techniques. During training, the model learns to predict the next word in a sentence based on the context provided by the preceding words. This process, known as autoregressive language modeling [16], enables the model to capture complex language patterns and semantics. Subsequently, the base model can undergo a fine-tuning process with human feedback and additional refinement through reinforcement learning, guided by a reward model trained using supervised methods. However, while general-purpose LLMs, whether closed-source (e.g. ChatGPT) or open-source (e.g. Llama2 [17,18]), have advanced clinical term identification tasks, they are known to fabricate or “hallucinate” citations, references, and source links [19]. This limitation restricts their suitability for concept normalization.

Recent studies have provided compelling evidence that the fine-tuning of LLMs with specialized medical data sources can facilitate their adaptation to specific tasks within clinical settings [20-22]. Fine-tuning is a process in which an unsupervised pre-trained LLM is further trained on a smaller, task-specific dataset to adapt its parameters to a specific task. This process involves updating the weights of the model’s layers using the task-specific data while retaining the knowledge learned during pre-training, enabling the model to better perform on the target task. For instance, Yang et al. successfully developed an LLM model by fine-tuning BERT and GPT to extract and recognize HPO phenotypes in clinical texts within the presence of non-HPO phenotypes, typos, and semantic differences with the model’s original training data [23]. However, that study did not consider the concept normalization task. In our study, we hypothesize that by fine-tuning LLMs using rare-disease-specific corpora and terminologies or ontologies, we can significantly augment their capacity to handle synonyms and textual variations, thereby enabling them to more precisely capture the intricate nuances woven into clinical texts. Consequently, the fine-tuned model has the potential to offer a nonstop solution for the critical task of recognizing standardized concepts from clinical narratives, an imperative need in the field of rare disease patient phenotyping.

## METHODS

### Overview

**Figure 1** provides an overview of the study design. We hypothesize that fine-tuned LLMs using a vocabulary-derived corpus will help overcome the challenge of clinical concept normalization. The pretrained model we used in this study is Llama 2 [18], an open-source LLM developed by Meta, which utilizes a transformer architecture similar to GPT models but is optimized for better parameter efficiency. We fine-tuned the Llama2-7B model, comprising 7 billion parameters, using generated sentences incorporating clinical concepts sourced from the HPO vocabulary (detailed in Data Source). In contrast to instruction fine-tuning, the Llama 2 model operates by completing a user input. For example, giving Llama 2 the prompt “The color of an apple is: ” yields an output of “red”. We adopted this element when fine-tuning and evaluating our model. In total, we fine-tuned two Llama2-HPO-Normalization models. The initial NAME model was fine-tuned using only standard HPO concept names without providing synonyms. The second NAME+SYN model was fine-tuned using standard concept names with half of each concept’s associated synonyms. For example, the concept “Hearing impairment” (HP:0000365) has six synonyms (“Deafness”, “Hearing defect”, “Hearing impairment”, “Hypacusis”, “Hearing loss”, “Hypoacusis”), but we only used three of the six to fine-tune the model. We assessed each model’s performance by constructing various prompts with different phenotype terms, including standard concept names, concept names with spelling errors, synonyms listed in the vocabularies (but not used in the fine-tuning), relevant terms cross-referenced from other vocabularies (e.g. SNOMED-CT) and laymen’s terms generated by ChatGPT. The performance is measured as the ratio of prompts that models can identify the correct HPO IDs for the phenotype terms (detailed in Evaluation). We used the Llama2 base model, GPT-3.5, and an index-based approach as benchmarks to evaluate the performance of our fine-tuned models.

**Figure 1.**
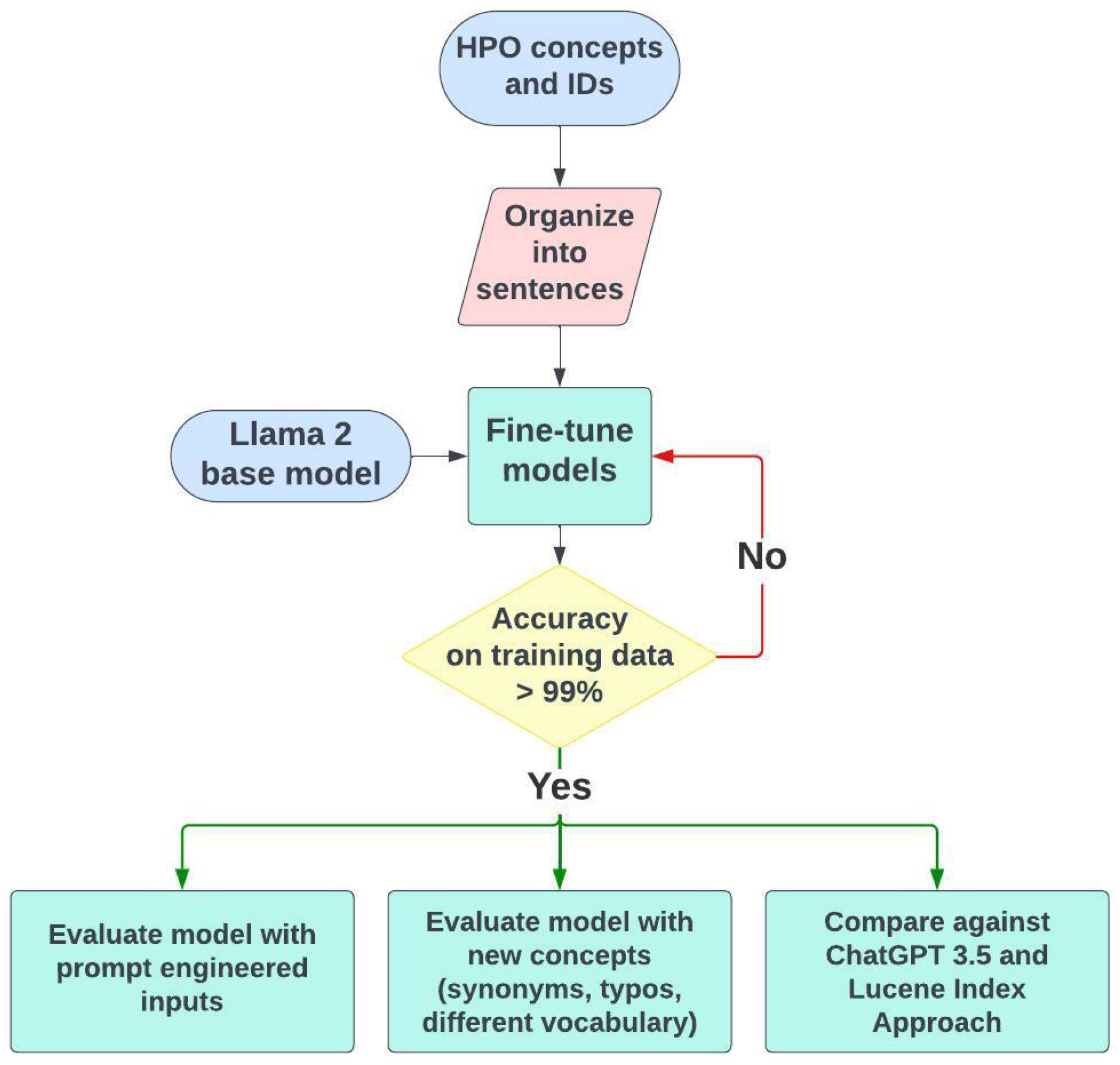
Overview of the study design.

### Data Source

We used a template-based approach to generate sentences used for fine-tuning Llama2. The NAME corpus consisted of sentences generated by associating each concept’s ID and only its standard concept name. The sentences we used for fine-tuning process are derived from this template: “The Human Phenotype Ontology term [CONCEPT] is identified by the HPO ID [HP ID].” An example is “The Human Phenotype Ontology term Hearing impairment is identified by the HPO ID HP:0000365”. The sentence has decent amounts of textual context with the full spelling of HPO and providing easy instructions for the model to correlate an HPO term to its identification tag. Furthermore, we constructed NAME+SYN corpus that consisted of sentences generated by both standard concept names and half of their synonyms (as annotated in the vocabulary). The HPO vocabulary consists of 17,066 distinct phenotype concepts. Therefore, the NAME model was fine-tuned with 17,066 distinct terms (standardized names), and the NAME+SYN model was fine-tuned with 31,737 terms (names + half of synonyms).

### Fine-tuning strategy

We utilized an autoregressive objective to fine-tune the normalization models as the next token prediction task. The autoregressive objective is a key concept in sequence modeling, where the goal is to predict the next element in a sequence based on previous elements. Mathematically, it involves maximizing the likelihood of observing the next element given the model’s predictions so far:

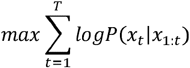

where *x*_*t*_ is the current token in the sequence and *x*_1:*t*_ is the sequence of previous tokens.

The fine-tuning was conducted on 4 NVIDIA A100 GPUs, with significant speed-up through low-rank adaptation (LoRA) [24]. The parameters *θ* were updated via learning a low-rank transformation matrix *W* using a small amount of task-specific data:

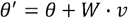

Where *v* is a vector obtained by aggregating information from the task-specific dataset. Earlier variants of the model underwent ten epochs to evaluate if the fine-tuning was functioning properly. Once the fine-tuning was confirmed to work, the number of training epochs gradually increased to assess how the model performs after more training. The data used to train the model, including the number of sentence variations and clinical concepts, also increased once the simplistic prototype models achieved functional results. The final training epoch was set as 100 as we observed that the training loss values became flat by 100 epochs. The fine-tuning process is implemented using the transformers and datasets module developed by HuggingFace [25]. Default parameters were used, except that the r value is set to 32 (so that 0.248% of the Llama 2 parameters are trainable in LoRA model) and that the batch size is changed to 128 to fit the GPU memory.

### Evaluation of the models

We assessed the performance of the models when presented with varied prompts and terms. The first part of the evaluation involved testing the models against different inputs via prompt engineering. Prompt engineering maintains the same phenotype terms as used in the training data but changes the query sentence structure (e.g. “HPO ID of [query term] is.”). This allows us to evaluate how well the models performed given “foreign” prompts (i.e. sentence not seen in the training data) with the same query terms.

The second part of the evaluation assesses the model’s adaptability to alterations in phenotype terms to which they have not been fine-tuned previously. The evaluation prompt “The Human Phenotype Ontology term [query term*] is identified by the HPO ID” maintains the same sentence structure but uses different input [query term*] than the fine-tuned corpus. The modified query terms can fall into one of the following categories: (1) standard names (as seen in the training set); (2) standard names with typos; (3) synonyms not seen in the fine-tuning set; (4) associated terms found in another vocabulary such as SNOMED-CT [26]; and (5) laymen’s term generated via ChatGPT 3.5. Synonyms defined in (3) and (4) were sourced from a list of concept synonyms provided by the HPO annotation database. For example, synonyms of “Hypoplastic hippocampus” include “Small hippocampus” and “Undeveloped hippocampus”; all three terms correlate to the HP:0025517 but differ textually. For ChatGPT, we used the prompt “Please generate five synonyms for the given phenotype term. For example, if the phenotype term is “Loss of consciousness”, return [“Fainting”, “Loss of consciousness”, “Passing out”]. Phenotype term: [HPO name]” to generate 500 terms. All synonyms used during evaluation were not included in the fine-tuning process. Typos include simple and complex typos. Simple typos were introduced randomly by altering one character from the original concept name based on keyboard proximity; for example, ‘i’ can be changed to ‘u’, ‘j’, ‘k’ and ‘o’. Complex typos were implemented by randomly altering 20% of characters (up to three) in a term. This enables us to more effectively assess models’ practical utility in real-world applications, where typos, synonyms, and laymen’s descriptions are commonly encountered.

We compared the performance of our models with three benchmarks: (1) concept IDs identified via an indexing-based information retrieval approach, (2) GPT-3.5 normalized concept IDs, and (3) pre-trained Llama 2 base model normalized concept IDs. For indexing-based approach, we indexed all standard concept names and synonyms defined in the vocabularies, with Lucene-based technology[27] and the Python Woosh library. The BM25 algorithm[28] was employed for information retrieval when a “query” term was supplied. The top-1 ranked concept ID (if returned) was retrieved as the normalized concept ID. We implemented two approaches for concept retrieval: ‘AND,’ which requires all tokens in the query term to be found in a returned concept, and ‘OR,’ which requires at least one of the tokens to be matched. For both ChatGPT and Llama 2, we used the prompt “The Human Phenotype Ontology term [query term] is identified by the HPO ID” for concept ID normalization.

## RESULTS

### Performance on the Llama 2 Base Model

Before we began the fine-tuning procedure, we assessed the performance of the Llama 2 base model. The Llama 2 base model is unable to associate HPO terms with their respective IDs (Figure 2). For example, when inputted with concept normalization prompts such as “The Human Phenotype Ontology term Vascular Dilatation is identified by the HPO ID,” the model outputted an arbitrary string of numbers unrelated to the HPO ID.

**Figure 2.**
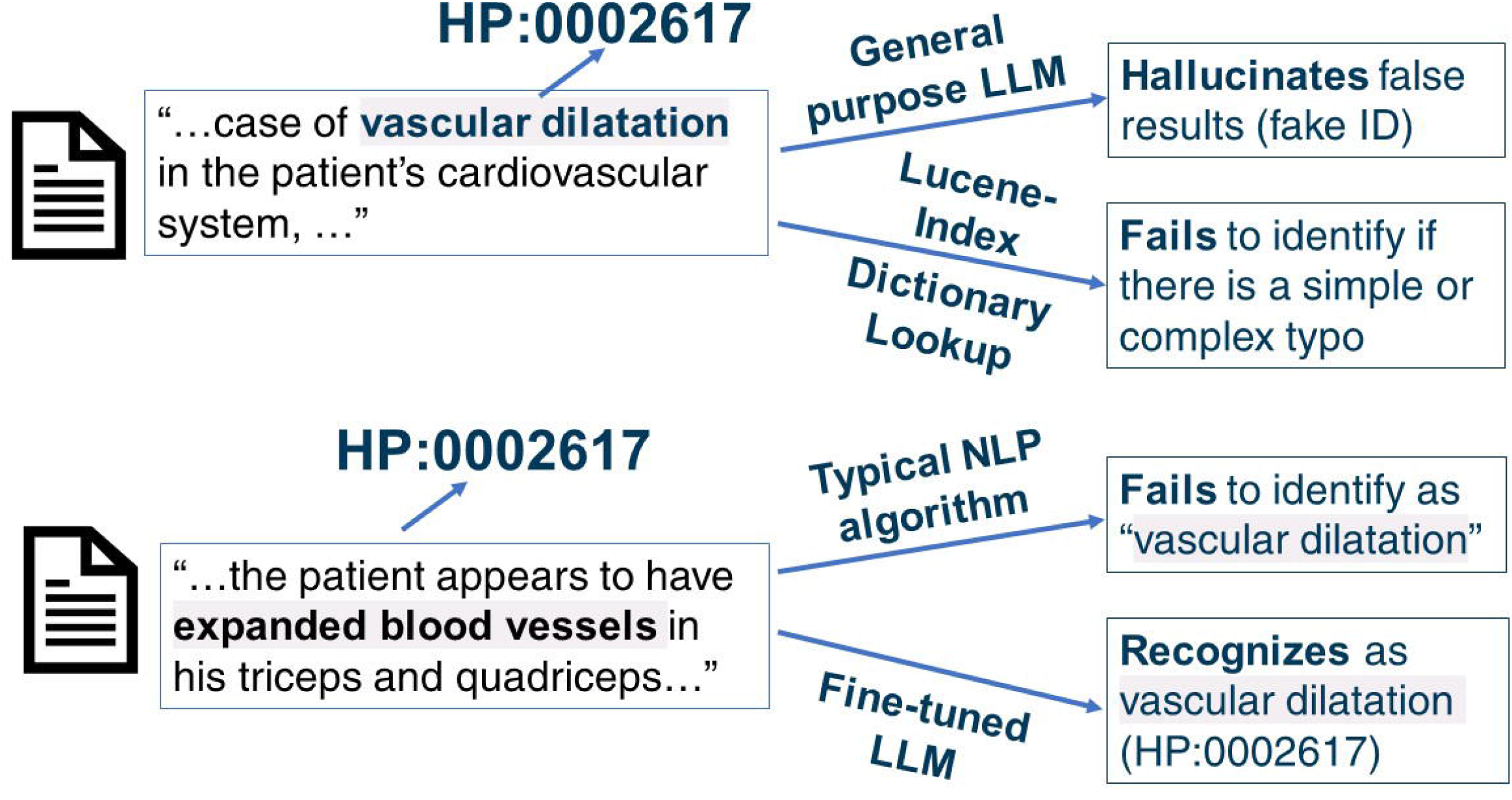
Examples of the ineffectiveness of traditional approaches and general-purpose LLMs (e.g. ChatGPT) at clinical concept normalization.

### The performance of fine-tuned models

Using Llama 2 base model with 7 billion parameters, we generated training sentences incorporating clinical concepts sourced from the HPO vocabulary and fine-tuned two HPO normalization models. The NAME model was fine-tuned using only standard HPO concept names, while the NAME+SYN model was fine-tuned using standard concept names with half of each concept’s associated synonyms. Following 100 epochs of fine-tuning, both models achieved nearly perfect accuracies when prompted with the original training data. In many incorrect cases, the inputted HPO term names were often lengthy and contained commas such as “Low-set, posteriorly rotated ears.” We suspect that this type of complex and long input could have confused the model and is the reason behind its incorrect identification. When different sentence structures were used (with the same query term as seen in the training corpus), such as “HPO ID of [query term] is,” the models struggled to normalize the terms. The model’s decreased performance against different types of sentence structures is expected given that the fine-tuning data includes one type of sentence structure.

### The performance on input with typos

When introducing typos into concept names, the performance decreased significantly in both HPO concept normalization models. We first tested simple typos of single-character modifications to the name. The replacement characters were implemented based on their proximity to other characters on the keyboard. For example, potential typo candidates for the letter ‘m’ include ‘n’, ‘j’, and ‘k’. The NAME model identified misspelled concept IDs with a 10.2% correction rate while the NAME+SYN model demonstrated a 36.2% accuracy (**Table 1**). For example, the models performed poorly in terms of typos such as “Bascular dilatation” and “Aneyrysms” instead of “Vascular dilatation” and “Aneurysms” respectively. However, when we include 30 additional fine-tuning epochs by altering the training sentences to include randomly generated typos, the performance for NAME+SYN model increased to 61.8%. For comparison, the indexing-based concept normalization Lucene-Index “OR” approach achieved a 45% accuracy rate, while the Lucene-Index “AND” approach can fail in nearly all cases, with a rate of 1.5%. We further tested the models with complex typos by introducing 3 altered characters into the input term. The performance dropped dramatically. The NAME model correctly identified complex typos with a 10.2% accuracy, and the NAME+SYN model performed with an accuracy of 8.2%. Similarly, when we include 30 additional fine-tuning epochs by altering the training sentences to include randomly generated single-character typos, the performance for NAME+SYN model increased to 25.3%. The indexing-based concept normalization using “OR” achieved an 18.9% accuracy whereas the ‘AND’ approach failed completely once more, with a rate of 0.7%. Upon examining the typos, we found them to be very complex, and often difficult for humans to accurately identify.

**Table 1.**
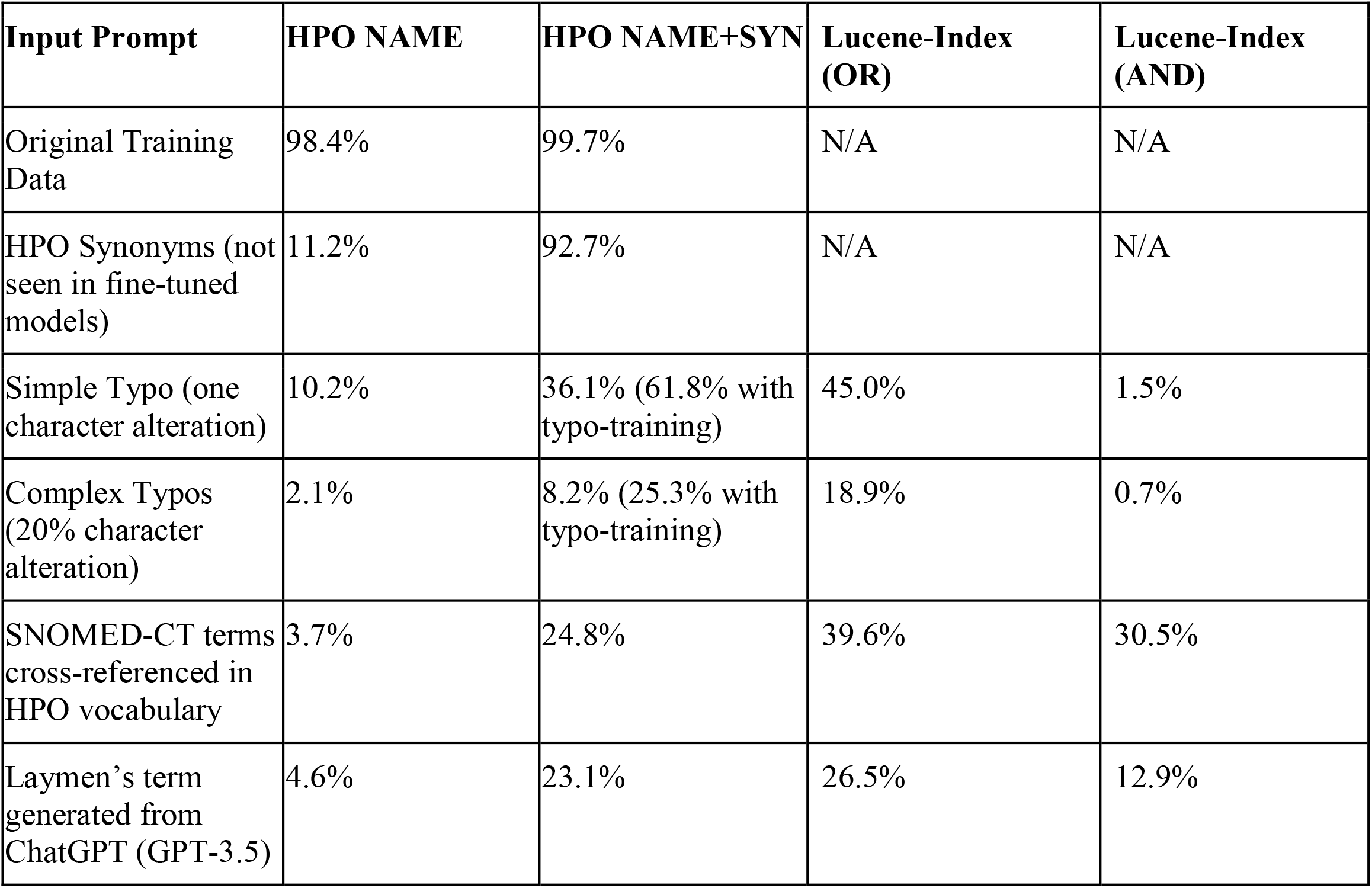
Accuracies of the two fine-tuned Llama models (HPO NAME and HPO NAME+SYN) and two Lucene-Index methods when prompted with various query terms.

### The performance on synonyms and laymen’s terms

We further tested the models on query terms they had not fine-tuned on. For this analysis, we first used HPO synonyms as the testing data (on average, there are 2.7 HPO synonyms per HPO concept). The NAME model achieved 11.2% accuracy, likely due to limited variation for a single ID in the fine-tuning set (**Table 1**). In contrast, the NAME+SYN model, fine-tuned with half the HPO synonyms, performed much better at 92.7%. In addition, we tested models’ performance on synonyms cross-referenced from SNOMED-CT and 500 ChatGPT-generated laymen’s terms. For SNOMED-CT synonyms, the NAME model achieved an accuracy of 3.7% while the NAME+SYN model improved to 24.8%. Tested against the 500 ChatGPT-generated terms, the NAME and NAME+SYN models achieved accuracies of 4.6% and 23.1%, respectively. As a comparison, the indexing-based approach using “OR” achieved an accuracy of 39.6% (30.50% using “AND”) and 26.5% (12.9% using “AND”) in correctly normalizing SNOMED-CT synonyms and ChatGPT-generated terms, respectively.

### The performance of ChatGPT (GPT3.5)

Mainstream LLMs such as ChatGPT are excellent at name entity recognition tasks, but we wanted to analyze whether they also possess accurate concept normalization abilities. Using ChatGPT 3.5 (accessed in September 2023) as a benchmark, it correctly identified ∼20% of HPO terms’ IDs. The correctly identified concepts are relatively common in clinical notes like “Diabetes mellitus HP:0000819” and “Hypertension HP:0000822”. ChatGPT generally fails on less commonly seen phenotypic features: it either claimed unfamiliarity, insisted the term did not exist, or generated imaginary (non-existent) HPO IDs. For example, when tasked to identify the HPO ID of “Vascular dilatation,” ChatGPT does not recognize the term as of its update in 2022. However, ChatGPT suggests a non-existent, “Arterial dilation,” with HPO ID, HP:0012824, which corresponds to the HPO concept “Severity.” ChatGPT not only hallucinates HPO IDs but the entire HPO concept names themselves. The hallucinated HP IDs follow the same formatting as the standard HPO ID, but the actual ID itself is incorrect. In other incorrect cases, ChatGPT claims the specific HPO term provided does not have a corresponding HPO ID but has offshoots, such as “neoplasm” and “Abnormality of the upper arm.” Both of these examples have their respective HPO IDs but ChatGPT claims otherwise. Additionally, the offshoot HPO terms and IDs it provides are incorrect, similar to that observed in the case with “arterial dilation” noted above. Our benchmark of ChatGPT indicates that mainstream LLMs fail at clinical concept normalization tasks. However, we acknowledge that ChatGPT is constantly being updated, and it is likely that more recent versions of ChatGPT can correctly identify more HPO terms.

### Illustration of End-to-End Model

To illustrate how the proposed approach can be used in an end-to-end setting to facilitate genetic diagnosis of rare diseases, we illustrate an example of using a public, de-identified clinical note on a patient with idiopathic progressive cognitive decline and other phenotypic features, previously reported [29](**Figure 3**). In a two-step approach (shown as in **Figure 3A**), various concept extraction tools, including traditional named entity recognition tools, ChatGPT, or more recent GPT-based phenotype extraction tools [23] can extract specific mentions of phenotypes from clinical notes first. And then these phenotype mentions are then normalized into HPO concepts with the corresponding HPO IDs using either our fine-tuned models or indexing-based approach. These HPO IDs can be used in software tools such as Phenomizer [30] and Phen2Gene [31] to prioritize candidate diseases in combination with genome or exome sequencing data.

**Figure 3.**
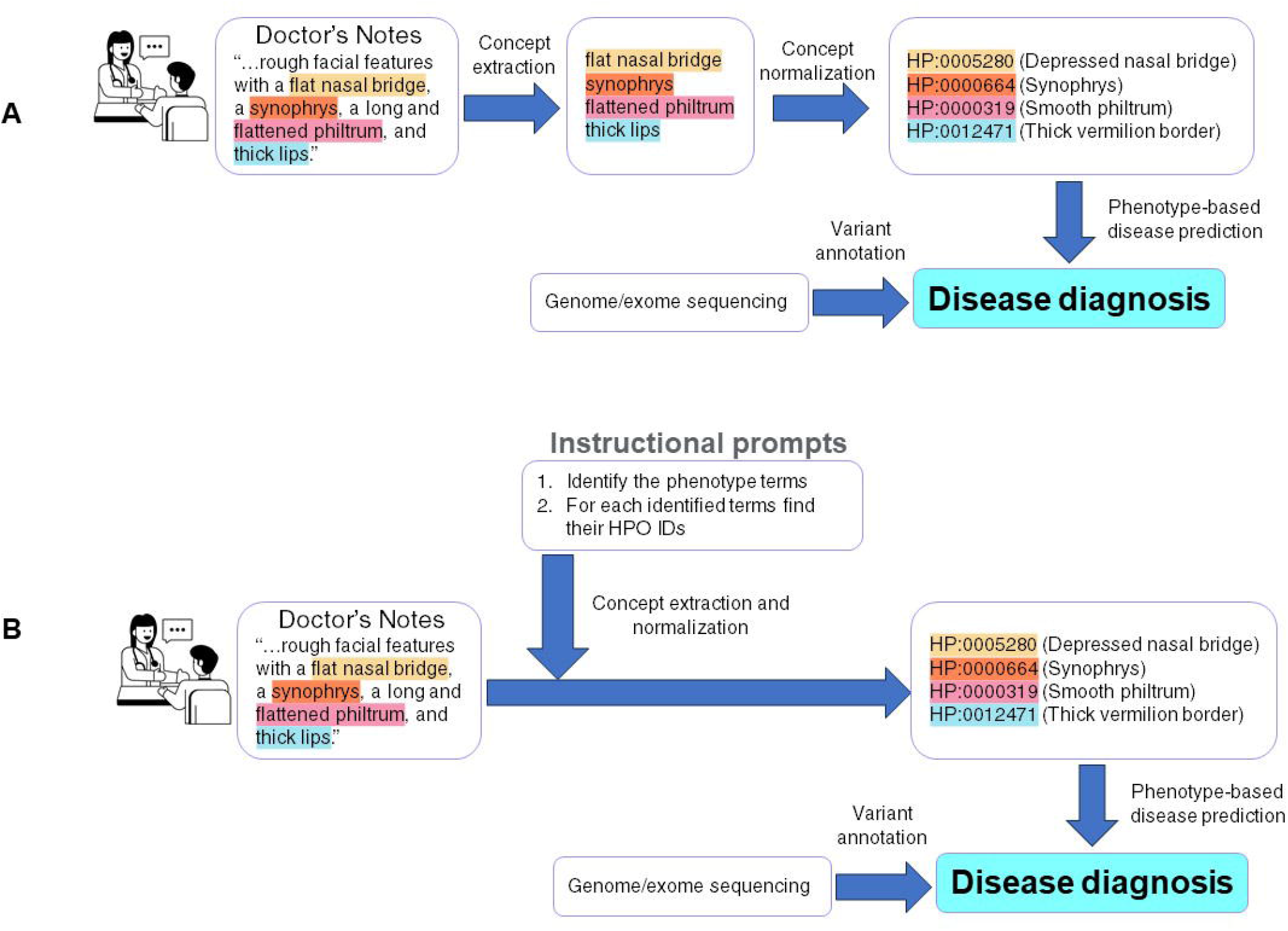
A case study showing how a concept normalization tool can be used in an end-to-end workflow to facilitate the prioritization of diseases and the phenotype-driven analysis of genome/exome sequencing data. (A) Two-step approach; (B) One-step approach.

However, one of the benefits of our fine-tuned approach is it can be used to develop a specific ChatBot like LLM (e.g. Llama2-Chat[32]) and use a “instruction prompt” to provide a one-step approach (as shown in **Figure 3B**).

## DISCUSSION

Compared to conventional national language processing algorithms, LLMs such as ChatGPT can effectively recognize synonyms of standardized vocabularies, thereby enhancing their efficacy in identifying phenotypes within clinical narratives. However, mainstream LLMs like ChatGPT fail at clinical concept normalization tasks. Our fine-tuning LLM helps to bridge this caveat in standardizing clinical concept normalization by accurately associating a clinical concept’s name directly with its respective identifier in ontologies (e.g. HPO). Our fine-tuned models demonstrated robustness to various prompts and different query terms, handling challenges such as misspellings, synonyms, laymen’s terms, and concepts from other ontologies like SNOMED-CT. However, several issues emerged during our evaluation, prompting considerations of additional improvement.

Throughout the fine-tuning process, we produced multiple variants of our fine-tuned model, each with varying amounts of training epochs and data. Our first variants trained on roughly 20 epochs had much lower accuracies than our current models but performed better than the Llama 2 base model without fine-tuning. Generally, increasing the number of training epochs correlated with improved accuracy until 60-70 epochs. In earlier iterations, we fine-tuned the model with multiple training sentences (more than one template). However, this approach increased training time significantly while yielding little to no improvements in the concept normalization task.

In general, both models can process various inputs and demonstrate robust performance if the query term is included in the fine-tuning corpora. The model, however, had a subpar performance when tasked with input sentences with diminishing amounts of textual context, suggesting more context (similar to the training sentence) in the input results in higher accuracies. Comparing the fine-tuned LLM to the indexing-based approach using ‘OR,’ the LLM did not perform better. However, many concept mapping implementations, such as the search function in the HPO.jax website, rely on the ‘AND’ approach for efficient consideration. In this context, the fine-tuned LLM outperforms the ‘AND’ based indexing approach. Importantly, the LLM-based concept normalization approach can seamlessly integrate with LLM-based concept recognition tasks, providing a more straightforward solution. Looking ahead, the use of larger models (such as 70B or even larger models like GPT-4) should be considered to further enhance performance.

The inaccurate results from our fine-tuned models were typically off by one or two digits from the end of the ID compared to the correct answer, indicating the model is close to associating those terms with their identifiers. This observation may be linked to the organizational structure of ontology IDs. Concepts with only the last digits differing often share the same parents in the concept hierarchy. This semantic closeness between two IDs could potentially contribute to errors in the ID identification task. Additionally, we noticed that the IDs were tokenized into digit-sized segments. This observation could explain the “last-digit” error, as LLMs ultimately aim to predict the next tokens. An alternative approach is to enhance fine-tuning by creating a customized tokenizer that treats the entire ID (e.g., HP:0004413) as a single token, rather than breaking it up into individual characters (e.g., “HP:”, “0”, “0”, “0”, “4”, “4”, “1”, “3”). This modification can potentially enable the model to capture more nuanced semantic relationships between concept names and their corresponding IDs.

Regarding the models’ performances against synonyms, instances of incorrect answers from the model often stemmed from inaccuracies related to n-gram concepts with special tokens such as parentheses and hyphens. Since almost none of the fine-tuning data included hyphens, it suggests a potential reason why the models performed poorly when handling terms with hyphens. This might also explain why SNOMED-CT synonyms presented challenges for both models since a majority of the SNOMED-CT synonyms include a hyphen. For example, the SNOMED-CT synonym of “Abnormality of the kidney” includes “Kidney - Abnormal” and “Kidney structure - Defect”, both of which have the hyphen as key to the concept meaning. Upon removing hyphens from each term, the models saw a ∼5% increase in normalization accuracy.

Another source of error is due to certain HPO concepts and their respective synonyms having different semantic meanings, especially without the clinical context. For example, “HA” can be abbreviated to “Headache”, but can also be a representative of “ Hemolytic Anemia”. Deciphering the abbreviation is straightforward within context, but a lack of background makes this term ambiguous. Given that the prompts evaluated in this study lack clinical context, future efforts should focus on constructing prompts using clinical narratives. This will help assess whether abbreviations can be accurately normalized within a more realistic clinical setting and eventually provide a single prompt solution for both entity recognition and concept normalization.

## CONCLUSION

Our fine-tuned Llama 2 model further advances the concept normalization task by linking identified phenotype terms with their respective identifiers. This approach can be extended to other medical language processing tasks, normalizing recognized medical entities to standard medical concepts in a controlled vocabulary. In a clinical setting, standardized phenotypic concepts can be used by many other informatics tools to identify disease-causal variants, rank candidate diseases, and forecast disease risk, thereby improving diagnostic and treatment accuracies. We plan to enhance the model by incorporating more data such as genes, drugs, and phenotypes, to standardize vast amounts of information. The model has the potential to generate knowledge graphs from narratives by linking diseases, phenotypes, genes, and drugs in a standard manner, therefore revealing previously unestablished relationships and outcomes.

## Contributorship Statement

A AW developed the software code, performed the computational experiments, analyzed the data, and wrote the manuscript. CL guided the interpretation of results and generated simulation data. JY advised on programming and model selection. CW conceived and supervised the study. All authors read and approved the manuscript.

## Acknowledgments

We extend our gratitude to Dr. Herbert Chase for his mentorship in the Columbia University Summer Research Program.

## Data Availability

The software code and the fine-tuned Llama2 models (as asset files in software release) are available on GitHub (https://github.com/andywang-25/Llama2-HPO-Normalization).

